# Discrete and continuum approximations for collective cell migration in a scratch assay with cell size dynamics

**DOI:** 10.1101/219204

**Authors:** Oleksii M Matsiaka, Catherine J Penington, Ruth E Baker, Matthew J Simpson

## Abstract

Scratch assays are routinely used to study the collective spreading of cell populations. In general, the rate at which a population of cells spreads is driven by the combined effects of cell migration and proliferation. To examine the effects of cell migration separately from the effects of cell proliferation, scratch assays are often performed after treating the cells with a drug that inhibits proliferation. Mitomycin-C is a drug that is commonly used to suppress cell proliferation in this context. However, in addition to suppressing cell proliferation, Mitomycin-C also causes cells to change size during the experiment, as each cell in the population approximately doubles in size as a result of treatment. Therefore, to describe a scratch assay that incorporates the effects of cell-to-cell crowding, cell-to-cell adhesion, and dynamic changes in cell size, we present a new stochastic model that incorporates these mechanisms. Our agent-based stochastic model takes the form of a system of Langevin equations that is the system of stochastic differential equations governing the evolution of the population of agents. We incorporate a time-dependent interaction force that is used to mimic the dynamic increase in size of the agents. To provide a mathematical description of the average behaviour of the stochastic model we present continuum limit descriptions using both a standard mean-field approximation, and a more sophisticated moment dynamics approximation that accounts for the density of agents and density of pairs of agents in the stochastic model. Comparing the accuracy of the two continuum descriptions for a typical scratch assay geometry shows that the incorporation of agent growth in the system is associated with a decrease in accuracy of the standard mean-field description. In contrast, the moment dynamics description provides a more accurate prediction of the evolution of the scratch assay when the increase in size of individual agents is included in the model.

## 1 Introduction

*In vitro* cell biology assays are used to study the invasive properties of malignant cells, to quantify different mechanisms of wound repair, as well as in the discovery of potential drugs (Riss 2005; Edmondson et al. 2014; Shah et al. 2016). Typically, cells are placed on a two-dimensional substrate, and are allowed to migrate, proliferate, and interact with each other, as illustrated in Figure 1a-b. Different experimental geometries, such as circular barrier assays, are possible, and modern imaging technologies provide means of collecting high resolution images of the cell population as it evolves (Johnston et al. 2015).

An example of a two-dimensional cell biology assay is given in Figure 1a-b. This kind of experimental design is routinely referred to as a *scratch assay*. Scratch assays are initiated by uniformly distributing a population of cells on a cell culture plate, which is then incubated for some time to allow cells to attach to the substrate and for the density of the monolayer of cells to increase. After incubation, a sharp-tipped instrument is used to scratch the monolayer to produce an artificial wound (Liang et al. 2007; Jin et al. 2016; Grada et al. 2017). The rate of the recolonisation of the wound space is then observed over time and has been demonstrated to depend on the rate of cell motility, the rate of cell proliferation, and the strength of cell-to-cell interaction forces. It is well known that quantifying the roles of these different mechanisms is challenging, as similar population-level outcomes can arise from different relative contributions of these separate mechanisms (Treloar et al. 2013). One way of overcoming these issues is to modify the experimental procedure to deliberately separate the effects of cell migration from the effects of cell proliferation, and this approach is routinely used to improve our understanding of the role of cell motility in wound healing and malignant spreading (Glenn et al. 2016; Nyegaard et al. 2016; Grada et al. 2017). The experimental images in Figure 1c-d show a scratch assay that is prepared in exactly the same way as the experimental images in Figure 1a-b except that the cells are treated with a drug to inhibit proliferation (Shah et al. 2016). A visual comparison of the images in Figure 1a-b and Figure 1c-d shows that the combined effects of cell migration and cell proliferation lead to a more rapid wound closure. The experiment in Figure 1c-d involves treating the cells with a drug called Mitomycin-C (Sadeghi et al. 1998). Mitomycin-C is a chemotherapy drug that suppresses mitosis by blocking DNA replication (Sadeghi et al. 1998). While Mitomycin-C is known to prevent cell proliferation without inhibiting cell migration (Simpson et al. 2013), a consequence of treating cells with Mitomycin-C is that cells increase in size, as if they are about to divide into two daughter cells, but the process of division does not take place. Therefore, cells that are treated with Mitomycin-C do not divide, but instead they approximately double in size over a period of approximately 24 hours, as illustrated in Figure 1e-f, where we show fibroblast cells in a circular barrier assay. The images in Figure 1e-f include a nuclear stain in red, and it is clear that the Mitomycin-C treated cells approximately double in size during the experiment. This change in cell size is typically neglected in mathematical models that describe the collective spreading of cell populations (Simpson et al. 2013; Jin et al. 2016).

**Fig. 1.**
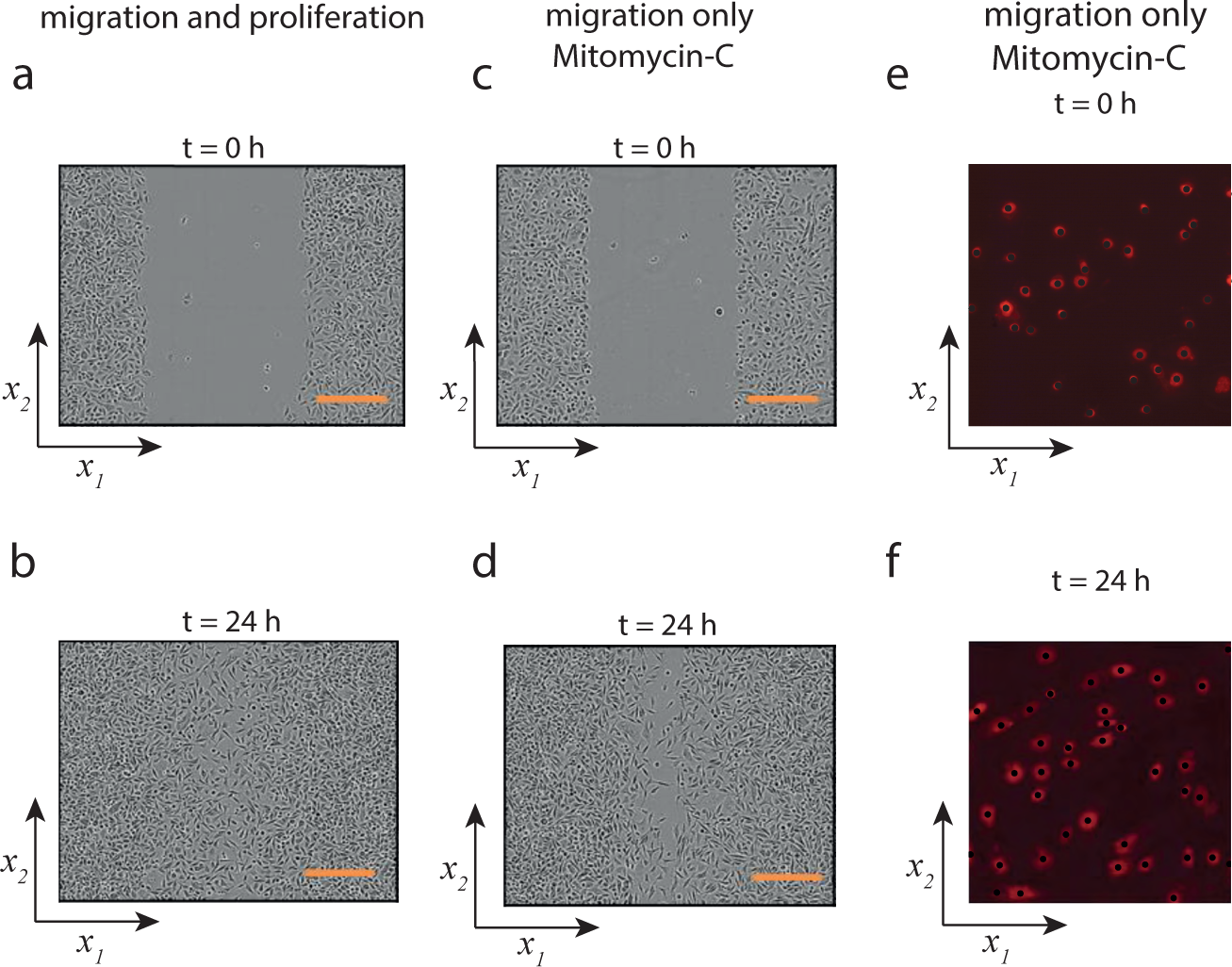
(a)-(b) An example of a typical scratch assay, where cells are allowed to close an artificially created gap. In this experiment the cells are PC-3 prostate cancer cells (Kaighn et al. 1979). (c)-(d) Scratch assay with PC-3 cells pretreated with Mitomycin-C to prevent proliferation. In (a)-(d) the scale bars correspond to 300 *μ*m. (e)-(f) Images showing individual 3T3 fibroblast cells in a circular barrier assay where the cells are treated with Mitomycin-C. In these images a cell nucleus stain is used, and each individual cell is superimposed with a black disk. We denote the two-dimensional coordinates as {*x*_1_*, x*_2_}, as indicated. Images in (e)-(f) show a square region of length 400 *μ*m. The images in (a)-(d) are reproduced with permission from Springer (Shah et al. 2016). The images in (e)-(f) are reproduced with permission from The Royal Society (Simpson et al. 2013).

Many experimental images, such as the images in Figure 1c-f, demonstrate the potential for significant changes in cell size which may influence the behaviour of the entire cell population since crowding effects are thought to be important in two-dimensional cell biology assays (Simpson et al. 2013). Many classical continuum models of collective cell behaviour do not incorporate any measure of cell size (Maini et al. 2004; Sherratt and Murray 1990). To address this limitation, another common approach to model collective cell behaviour is to use lattice-based stochastic models, where individual agents are allowed to move on a discrete lattice (Penington et al. 2011; Markham et al. 2015). Lattice-based models are attractive because they are conceptually straightforward, computationally efficient, and produce time-lapse images that are similar to images obtained from experiments (Simpson et al. 2013). While lattice-based models typically associate the cell size with the lattice spacing (Simpson et al. 2013), modelling the collective behaviour of populations of cells involving cell-to-cell crowding effects and dynamic changes in the size of individual cells in this approach is not straightforward. In particular, in a lattice-based model it is difficult to represent the cell size as a continuous function of time (Binder and Simpson 2016). An alternative approach is to use a lattice-free stochastic model (Codling et al. 2008; Galle et al. 2005; Newman and Grima, 2004). Lattice-free models can be more computationally demanding than lattice-based models when dealing with crowding effects. However, lattice-free models are much more appealing than lattice-based models because agents in the simulation can assume a continuous size, which may be allowed to change dynamically.

While computational implementations of stochastic models are well suited to capturing individual-level details, experimental data is routinely presented in the form of population-level and tissue-level data. Consequently, it is convenient to have access to some continuum approximation to describe the collective behaviour associated with the stochastic simulations. The continuum description often takes the form of a partial differential equation (Penington et al. 2011; Dyson et al. 2012). Most continuum-based models used to describe collective behaviour of cell populations invoke the mean-field approximation (MFA) (Sherratt and Murray 1990; Maini et al. 2004; Penington et al. 2011; Simpson et al. 2013). In effect, the MFA amounts to assuming that the positions of individuals in the population are independent. This assumption is widely invoked, both implicitly (Sherratt and Murray 1990; Maini et al. 2004) and explicitly (Penington et al. 2011; Dyson et al. 2012; Simpson et al. 2013). However, since spatial structure, such as clustering and patchiness, is often observed experimentally, the MFA is not always appropriate. Clustering and patchiness are observed in a range of natural processes, including cell biology experiments (Steinberg 1996; Treloar et al. 2013) and ecology (Levin and Whitfield, 1994), therefore it is also of interest to derive continuum limit descriptions that avoid the MFA, where appropriate. To achieve this, in this work we describe a lattice-free model of the collective spreading of a population of cells that incorporates cell motility, dynamic cell size changes, crowding effects, and cell-to-cell adhesion. We derive a continuum description using the standard MFA, as well as introduce an alternative continuum description using a more sophisticated moment dynamics approach (Middleton et al., 2014). Moment dynamics approaches are often used to describe spatially correlated populations in ecological applications and in the study of epidemics (Bolker and Pacala 1997; Keeling et al. 1997; Sharkey et al. 2006; Sharkey et al. 2015). However, moment dynamics approaches are less common in the study of collective cell behaviour. Moment dynamics approaches can invoke many different approximations to account for spatial correlations (Murrell et al. 2004; House 2014; Plank and Law 2015; Binny et al. 2015; Binny et al. 2016a; Binny et al. 2016b), and in this work we use the Kirkwood superposition approximation (KSA) which was originally developed to describe the spatial arrangement of molecules in liquids (Kirkwood 1935; Singer 2004), and only much later adopted to describe the spatial arrangement of individual cells in the collective cell spreading (Baker and Simpson 2010; Middleton, et al. 2014).

In this manuscript we present discrete and continuum descriptions of collective cell behaviour formulated in both one and two dimensions. Two-dimensional models allow us to reproduce the dynamics of experiments such as those depicted in Figure 1, and are perfectly suited for visualisation of the experiments. However, as we demonstrate in the Supplementary Material document, there is little motivation to use two-dimensional descriptions for the scratch assay geometry because the agent density, on average, does not depend on the vertical coordinate.

We denote the position of an arbitrary point in the computational domain by the vector **u** = {*x*_1_, *x*_2_}. The positions of two other arbitrary points in the domain are given by the vectors **u**^′^ = {*y*_1_, *y*_2_} and **u**^′′^ = {*z*_1_, *z*_2_}, and so on. We utilise the notation *x*, *y*, and *z* to denote the positions of distinct points in the one-dimensional domain when introducing continuum descriptions in one dimension. The position of the *i*th agent on a two-dimensional domain is **u**^(*i*)^. The position of an agent *i* in the one-dimensional discrete simulations is given by *x*(*i*). This choice of notation allows us to distinguish between the positions of agents in the discrete simulations and the coordinates of fixed points in the continuum description. Furthermore, it is consistent with our previous work which does not involve dynamical changes in agent size (Matsiaka et al. 2017).

This manuscript is organised as follows. In Section 2.1 we describe a stochastic lattice-free model of collective cell migration. The model incorporates cell migration, cell crowding effects and cell-to-cell adhesion, and allows individual cells in the population to change size dynamically. In Section 2.2 we introduce two continuum descriptions of the stochastic model: (i) a mean-field based approximation; and (ii) a moment dynamics approximation, based on the KSA. In Section 3 we compare the averaged data obtained from repeated simulations of the stochastic model with numerical solutions of both continuum approximations. Finally, in Section 4 we summarise our findings and discuss the opportunities for further research.

## 2 Methods

### 2.1 Langevin stochastic model

In this section we describe the lattice-free stochastic model used to simulate the collective behaviour of a population of *N* agents. Many two-dimensional cell biology experiments, such as the scratch assays depicted in Figure 1a-d, can be described using a one-dimensional coordinate system because the density of cells is independent of the vertical coordinate (Johnston et al. 2015). Therefore, we focus our attention and discussion on the one-dimensional discrete model in the main manuscript. Additional results and discussion relating to justifying the use of a one-dimensional model to describe a two-dimensional cell culture experiment is presented in the Supplementary Material document.

In this work we denote the cell diameter using *δ*(*t*) > 0, and assume that the dynamic change in cell diameter is logistic,

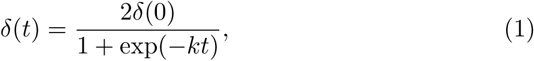

where the parameter *k* > 0 describes the growth rate, *δ*(0) is the initial cell diameter, and 
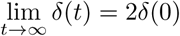
. Plots showing typical *δ*(*t*) for different choices of *k* are given in Figure 2. We note that the choice of using a logistic function for *δ*(*t*) is not essential, and all of the analysis presented here can be applied to any other suitable choice of growth model.

**Fig. 2.**
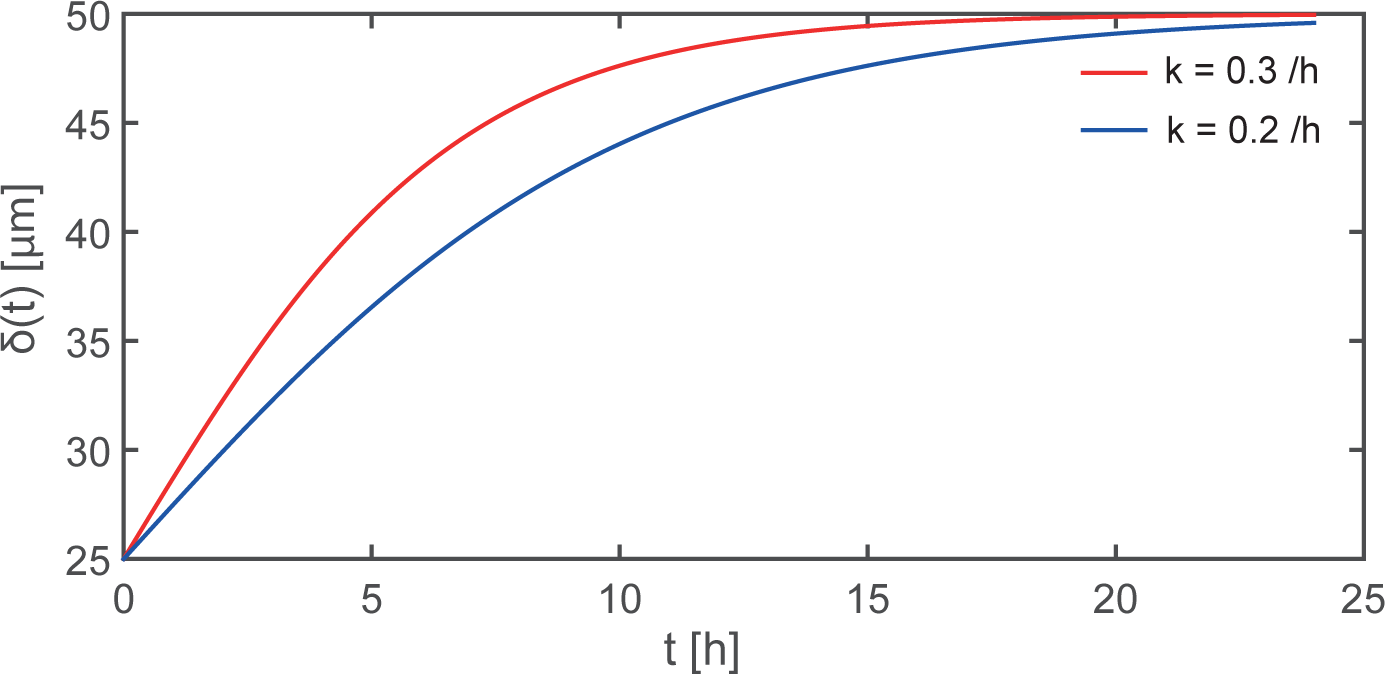
Logistic increase in agent diameter, given by Equation (1), for *k* = 0.3/h (red) and *k* = 0.2/h (blue). In both cases *δ*(0) = 25 *μ*m and 
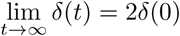
.

We assume that the movement of individual agents in the stochastic model is described by an equation of motion (Newman and Grima 2004; Middleton et al. 2014). We adopt the Langevin stochastic model where the collective behaviour of the population is described by a system of Langevin equations that can be written as

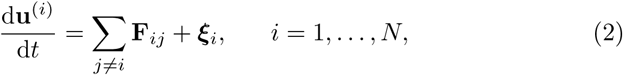

where **u**^(*i*)^ is the position of the *i*th agent in a two-dimensional space, **F***_ij_* is the interaction force between agent *i* and agent *j*, ***ξ****_i_* is the stochastic force acting on the *i*th agent, and *N* is the number of agents in the simulation.

The corresponding one-dimensional model is given by

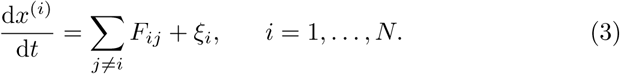

Since we consider unbiased movements of isolated individuals, we sample *ξ_i_* from a Gaussian distribution (Newman and Grima, 2004) with variance

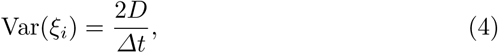

where *Δt* is the value of the time step used to solve the system of Langevin equations in a simulation of the stochastic model. Choosing *Δt* in this way ensures that the mean squared displacement of an isolated individual agent is independent of the time step chosen to simulate the stochastic model, and the mean squared displacement of an isolated agent is therefore 2*Dt*.

The force function, *F_ij_*, is chosen to be

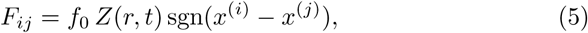

where *f*_0_ is a constant that describes the strength of the interaction forces, *Z*(*r, t*) is the dimensionless function describing how the interaction force depends on the separation of the agents, *r* = |*x*^(*i*)^ − *x*^(*j*)^|, *t* is time, and sgn is the *signum* function (Middleton et al. 2014; Matsiaka et al. 2017). A schematic showing the arrangement of agents in the model is given in Figure 3a-b, where we can see the effects of agent movement and the increase in the size of the agents with time.

We consider two main features of agent-to-agent interactions: (i) a short range repulsion force, which can be thought of as a resistance to deformations and crowding; and, (ii) longer range attraction forces which can be thought of as adhesion between agents. To model these two forces we use a modified Morse potential,

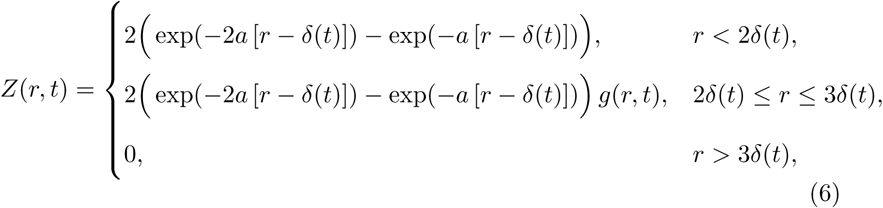

where *a* > 0 is a parameter that controls the shape of the force function, *δ*(*t*) is the time-dependent agent diameter, given by Equation (1) or some other appropriate functional form. Here, the spatial range of interactions is finite, and set to three agent diameters, giving *Z*(*r, t*) = 0 for *r* > 3*δ*(*t*). The function *g*(*r*) is the Tersoff cut-off function (Tersoff et al. 1987), which is included to capture the finite range of interactions,

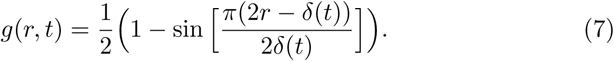

Figure 3c shows a typical interaction function, *Z*(*r, t*), over a period of 24 hours. At each instant in time the interaction force function incorporates repulsion at short distances, and attraction at longer distances, up to a finite range of three agent diameters. Since *δ*(*t*) increases with *t*, the interaction function also changes with time, and we can interpret the change in *Z*(*r, t*) with *t* in Figure 3c as a result of the increase in agent diameter with time. The choice of the force function in Equation (6) is not unique, but rather one of many other possible functional forms that incorporate short range repulsion and longer range attraction such as the Lennard-Jones potential, Hertz potential, or a nonlinear spring model (Byrne and Preziosi 2003; Jeon et al. 2010; Murray et al. 2012).

**Fig. 3.**
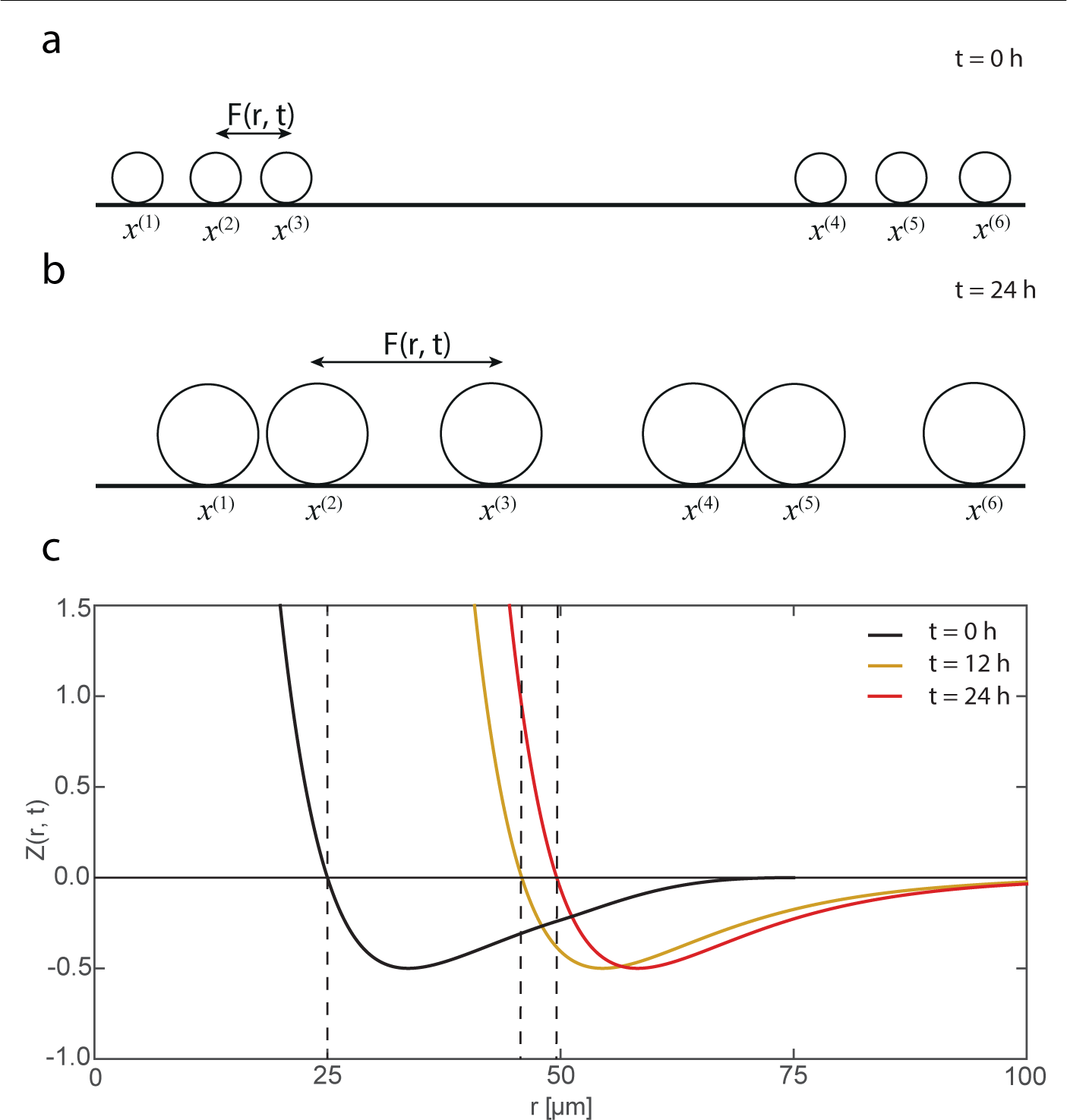
(a)-(b) Schematic illustration of individual agent motility and interaction forces in a population of agents where the diameters of individual agents double over an interval of approximately *t* = 24 h, as described by Equation (1). (c) Dimensionless potential, *Z*(*r, t*), at *t* = 0,12, and 24 h in black, orange and red, respectively. All plots correspond to *k* = 0.2 h and *δ*(0) = 25 *μ*m. The horizontal line at *Z*(*r, t*) = 0 denotes the change from short range repulsion (*Z*(*r, t*) > 0) to longer range attraction (*Z*(*r, t*) < 0). The series of three vertical dashed lines indicate the diameter of the agent at *t* = 0, 12 and 24 hours.

### 2.2 Continuum description

In this section we present two different continuum approximations of the lattice-free stochastic model described in Section 2.1. In particular, we present both a standard continuum approximation, based on invoking the MFA, and a moment dynamics continuum approximation, based on invoking the KSA. We note that the process of deriving both continuum approximations has been presented previously in the simpler case where the agent diameter is a constant (Middleton et al. 2014; Matsiaka et al. 2017). The focus of the current study is to consider the continuum approximations for the case where the agent diameter increases with time. Therefore, we do not repeat the derivation of the continuum limit descriptions here in the main document. Instead, complete details of the derivations are given in the Supplementary Material document, and here we focus on reporting the continuum descriptions and examining the accuracy of the continuum descriptions.

#### 2.2.1 Two-dimensional continuum model

Here we first present the two-dimensional continuum limit of the stochastic model, and later consider the one-dimensional analogue of this model. The mean-field continuum description is given by an integro-partial differential equation (IPDE),

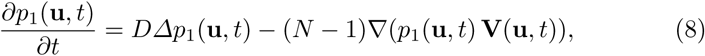

where the agent density, *p*_1_(**u**,*t*), depends on the position **u** = {*x*_1_, *x*_2_} and time *t*, *D* is the diffusivity, *N* is the number of agents, and

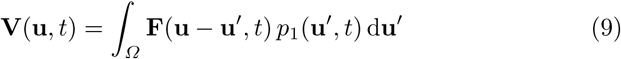

is the velocity field induced by the agent-to-agent interactions. The force function **F**(**u** *−* **u**^′^, *t*) is defined as

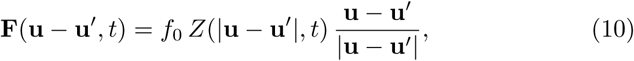

and depends on the separation distance, |**u** *−* **u**^′^| and time *t*.

The two-dimensional moment dynamics model, based on the KSA approximation (Singer 2004; Middleton et al. 2014), can be written as

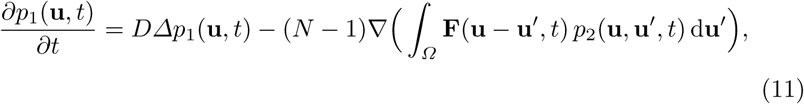

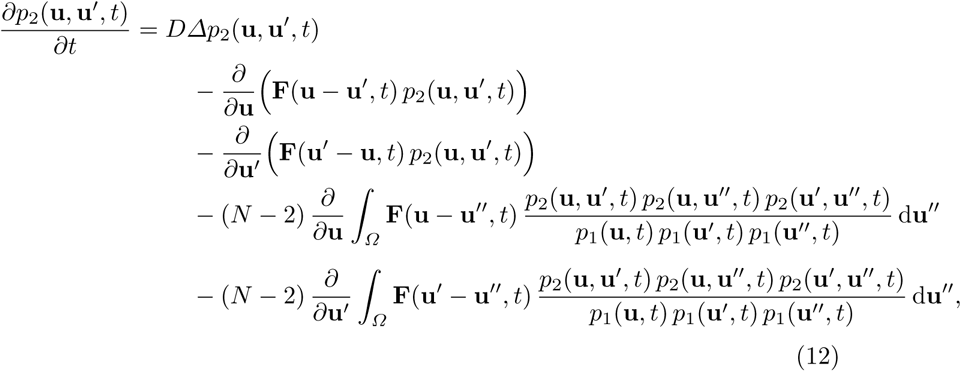

where *p*_2_(**u**, **u**^′^, *t*) is the density-density correlation function that captures correlations in the positions of agents at locations **u** and **u**^′^, at time *t* (Middleton et al. 2014).

#### 2.2.2 Simplified one-dimensional continuum model

The experimental images depicted in Figure 1a-d demonstrate the evolution of the cell population on a two-dimensional substrate. We note that the cell density in a scratch assay is independent of the vertical coordinate (Figure 1a-d) so that the experiment can be described in terms of a one-dimensional coordinate system. The underlying discrete model for Equation (8) and Equations (11)-(12) is a system of Langevin equations, Equation (3), as described in Section 2.1. To justify the one-dimensional approach we reproduce the experimental image in Figure 1(a) using two-dimensional Langevin model, and compare the density profiles obtained from the two-dimensional model with the results from the one-dimensional stochastic model (Supplementary Material document). Results presented in Figure 1 (Supplementary Material document) show that the simpler one-dimensional model produces similar results to the two-dimensional model for this special initial condition where the agent density is independent of the vertical coordinate. Motivated by these considerations, we neglect density variations in the vertical direction and write Equation (8) and Equations (11)-(12) in a one-dimensional format. We note that the use of one-dimensional continuum models to understand and interpret multidimensional transport phenomena with appropriate symmetry imposed by the initial conditions and boundary conditions is relatively common in both the mathematical biology literature (Callaghan et al. 2006; Khain et al. 2011; Smith et al. 2017), as well as in other areas of engineering and applied science (Simpson 2009).

The mean-field continuum limit of the one-dimensional stochastic model is given by an IDPE that can be written as

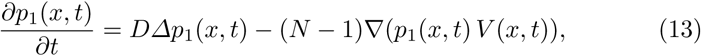

where the agent density, *p*_1_(*x, t*), depends on the position *x* and time *t*, and

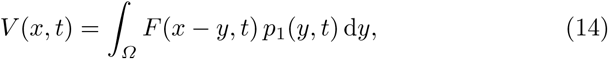

is the velocity field induced by the interactions between agents. We note that the diffusivity in Equation (13) is directly related to the stochastic force, *ξ_i_*, in the stochastic model, Equation (3).

As stated previously, continuum limit descriptions based on the MFA amount to assuming that the positions of agents are independent. However, in many practical situations, cells and other living organisms can adhere to each other and form clusters (Steinberg, 1996). In these situations the assumptions underpinning MFA-based continuum models are questionable. To address this limitation, we now make use of a more sophisticated moment dynamics continuum description that accounts for the density of agents and the density of pairs of agents. The moment dynamics model, based on the KSA, can be written as

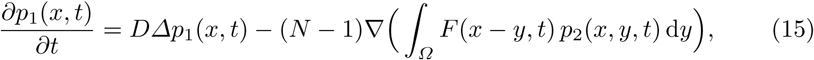

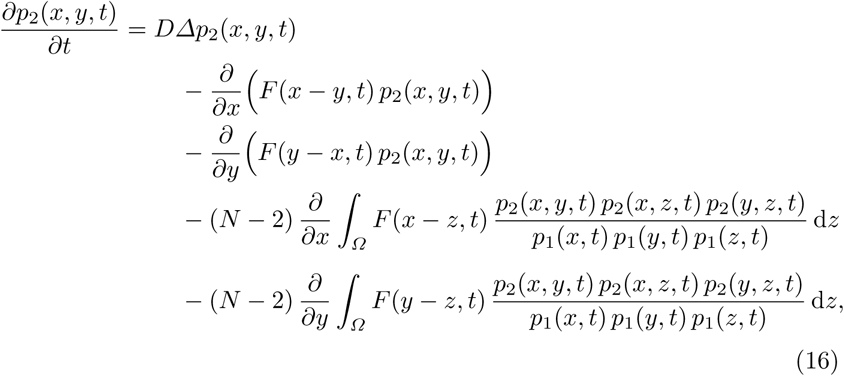

where *p*_1_(*x, t*) is the average density of agents at location *x* and time *t*, and *p*_2_(*x, y, t*) is the density-density correlation function that captures correlations in the positions of agents at locations *x* and *y* at time *t*.

## 3 Results and discussion

Here we focus our attention on a typical scratch assay geometry (Figure 1a-d). In this experiment the cell density does not depend, on average, on the vertical position (Figure 1a-d). Therefore, we can approximate this experiment as a one-dimensional problem.

The experimental images in Figure 1a-d show only a small region of the population, which extends well beyond the vertical boundaries of the image (Johnston et al. 2015). We apply periodic boundary conditions in all simulations (Middleton et al. 2014). The initial condition is given by sampling from a distribution, *α*(*x*), that is given by

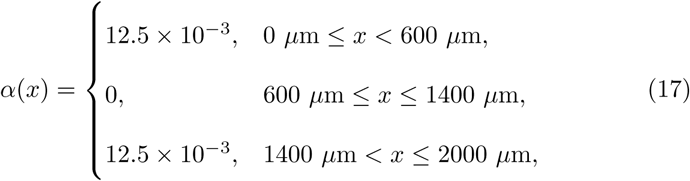

where the length of the domain, 2000 *μ*m, is a typical width of an experimental image (Jin et al. 2016). To initialise the stochastic simulations we randomly place agents in the two intervals, 0 ≤ *x* ≤ 600 *μ*m, and 1400 ≤ *x* ≤ 2000 *μ*m. This initial distribution of agents in the simulations mimics the initial distribution of cells in the images of scratch assays, given in Figure 1a and Figure 1c. In all of our results we report the agent density profiles in terms of both the dimensional density of agents, *p*_1_(*x, t*) [agents*/μ*m], as well as a non-dimensional agent density, 
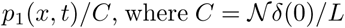
 is the carrying capacity density of agents with diameter *δ*(0). Here 
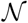
 is the maximum number of agents of diameter *δ*(0) that can be distributed, without compression, along a line of length *L*. In Equation (17) we choose the maximum density to be 12.5 × 10_−3_ cells/*μ*m because this corresponds to a non-dimensional density of approximately *p*_1_(*x, t*)/*C* = 0.625, and this is a typical initial density used in practice (Jin et al 2016; Liang et al. 2007). To initialise our simulations we place agents of diameter *δ*(0), at random, until the density of agents in the two intervals, 0 ≤ *x* ≤ 600 *μ*m, and 140 ≤ *x* ≤ 2000 *μ*m, is *p*_1_(*x, t*)/*C* = 0.625.

We describe the evolution of the scratch assay in three different ways. First, we perform individual realisations of the stochastic model to produce individual snapshots showing the distribution of agents in each simulation. Second, we perform a large number of identically prepared realisations of the stochastic model, and count the numbers of agents in *I* equally-spaced intervals across the domain. Averaging the number of agents in each interval allows us to quantify the spatial variation in the average agent density. Finally, we solve both the MFA- and KSA-based continuum models numerically, and compare the solutions of the continuum models with the average data from the suite of stochastic simulations to examine the accuracy of the two continuum descriptions. To quantify the accuracy of the two continuum descriptions we use

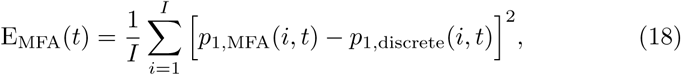

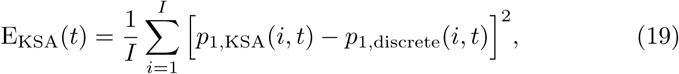

where, the index *i* denotes spatial node, *I* is the total number of nodes used to quantify the averaged agent density, *p*_1,MFA_(*i, t*) is the density of agents predicted by the MFA-based continuum model, *p*_1,KSA_(*i, t*) is the density of agents predicted by the KSA-based continuum model, and *p*_1,discrete_(*i, t*) is the density of agents predicted by averaging a suite of identically prepared realizations of the stochastic Langevin model. Here, E_MFA_(*t*) is a measure of the error associated with the MFA-based continuum description, and E_KSA_(*t*) is a measure of the error associated with the KSA-based continuum description.

We first examine the most straightforward situation where we consider a scratch assay with a population of cells where the diameter of cells remains constant. Following this preliminary case, we then examine two additional situations where we consider the same scratch assay except that the diameter of individual agents in the populations increases at different rates: (i) relatively slow growth, *k* = 0.2 /h; and (ii) faster growth, *k* = 0.3 /h. In all cases we fix the diffusivity to be a typical value for the PC-3 prostate cancer cell line, *D* = 1200 *μ*m^2^/h (Jin et al. 2016).

The stochastic model, Equation (3), is solved numerically using a first order explicit Euler method (Press et al. 2007). The number of individual realisations used to construct the density profiles is chosen to be 10^5^. This choice produces averaged density data with fluctuations that are two orders of magnitude smaller than the density data. For example, at *t* = 0, the density in the region *x* < 600 *μ*m is approximately 10*^−2^* agents/*μ*m, whereas the standard deviation of the agent density is approximately 10*^−4^* agents/*μ*m. After performing 10^5^ identically prepared simulations of the stochastic model, the spatio-temporal distribution of agent density is estimated by averaging results from identically-prepared realisations. The initial condition for the MFA-based continuum model, Equation (13), is *p*_1_(*x*, 0) = *α*(*x*), and the solution of the MFA-based continuum model is obtained by solving Equation (13) numerically, as described in the Supplementary Material document. Since we choose the positions of agents in the stochastic simulations to be random at *t* = 0, there are no correlations in the initial distribution of agents. Consequently, the initial condition for the KSA-based continuum model, Equations (15)-(16), is *p*_1_(*x,* 0) = *α*(*x*), and *p*_2_(*x, y,* 0) = *α*(*x*)*α*(*y*). To predict the evolution of the system using the KSA-based continuum model, numerical solutions of Equations (15)-(16) are obtained using techniques outlined in the Supplementary Material document. In all cases, the numerical solutions of both continuum models are obtained using a sufficiently fine spatial and temporal discretisation that the results are grid independent.

The results in Figures 4-6 compare solutions of the MFA- and KSA-based continuum models with the averaged results from stochastic simulations in the cases of: (i) no growth, *k* = 0 /h; (ii) relatively slow growth, *k* = 0.2 /h; and (iii) faster growth, *k* = 0.3 /h. To quantify the performance of the MFA and KSA models we compute the time evolution of E_MFA_(*t*) and E_KSA_(*t*), given by Equations (18)–(19), respectively. The only difference between the continuum-stochastic comparisons in Figures 4-6 is the rate of increase of the agent diameter, *k*. All other parameters in the simulations in Figures 4-6 are held constant to avoid ambiguity and to highlight the influence of agent growth on the dynamics of the population and the performance of the two different continuum descriptions.

The results in Figure 4a-c, Figure 5a-c and Figure 6a-c show stochastic simulations evolving from different realisations of the initial condition, Equation (17), to give a spatial distribution of agents after 24 and 48 hours. Note that the distributions of agents in Figure 4a-c, Figure 5a-c and Figure 6a-c are given as a series of 100 separate, one-dimensional simulations that are plotted adjacent to each other (Matsiaka et al. 2017). Presenting the stochastic results in this way is convenient because it highlights the randomness in the stochastic model. In general, we see that over a period of 48 hours the wound, of initial width 800 *μ*m, becomes recolonised by agents and the wound appears to close. Comparing the evolution of the stochastic models in Figure 4a-c, Figure 5a-c and Figure 6a-c, with the experimental images in Figure 1c-d suggests that this choice of parameters in the stochastic model is reasonable, as the rate of wound closure in the stochastic simulations is similar to the rate of wound closure in the experimental images.

In the case where there is no growth (Figure 4), the solution of the MFA-based continuum model matches the averaged agent density profile from the stochastic simulations very well (Figure 4d). Similarly, comparing the solution of the KSA-based continuum model with the averaged agent density profile (Figure 4e) reveals an excellent match. In this case, there seems to be littlejustification for use of the more complicated KSA-based continuum model as the simpler MFA-based model captures the evolution of the averaged agent density extremely well. Furthermore, quantitative comparison of the accuracy of the MFA-based continuum model with the accuracy of the KSA-based continuum model (Figure 4f) confirms that there is no advantage in using the KSA-based model for this problem where the size of the agents remains fixed.

In contrast, when we consider the situation where agents increase in size, *k* > 0 (Figures 5 and 6), the improved performance of the KSA-based continuum model becomes clear. Results in Figure 5d compare the evolution of the MFA-based model and averaged agent density data from the stochastic model, showing that there is a clear and visually discernible difference between two sets of profiles. This difference is quantified in Figure 5f. In contrast, the accuracy of the KSA-based continuum model, shown in Figure 5f, remains excellent. Similar comparisons between the performance of the MFA- and KSA-based continuum models in Figure 6 for faster growth confirms the improved accuracy of the KSA-based continuum model in the case when the agents are allowed to increase in size dynamically.

**Fig. 4.**
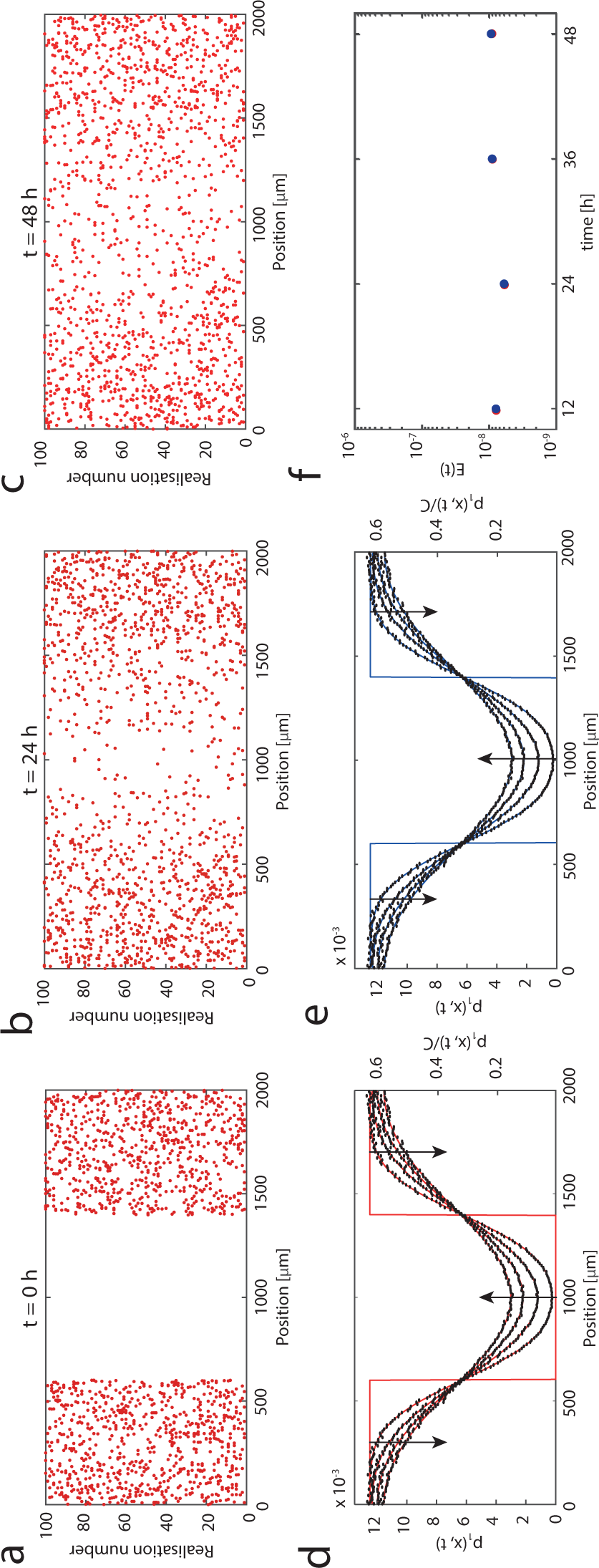
Comparison of the averaged agent density data from stochastic simulations and the solutions of the MFA- and KSA-based continuum models for a scratch assay with constant cell size, *δ*(*t*)≡25 *μ*m. Snapshots in (a)-(c) show 100 identically prepared realisations of the stochastic model, given by Equation (3), at *t* = 0, 24 and 48 h, respectively. The initial density is given by Equation (17). Results in (d) and (e) show the averaged agent density profiles using an ensemble of 10_5_ identically prepared simulations (black dots) with binsize 10 *μ*m. The averaged results are compared with solutions of the MFA-based continuum model, Equation (13) (red lines), and the KSA-based continuum model, Equation (16) (blue lines). Profiles are given at *t* = 0, 12, 24, 36 and 48 h, with the arrows indicating the direction of increasing *t*. Results in (f) show the evolution of the mean squared errors (Equations (18) and (19)) for the MFA- and KSA-based continuum models. Equation (3) is integrated with *Δt* = 2 × 10^−2^/h, Equation (13) is integrated with *Δx* = 4 *μ*m and *Δt* = 1 × 10^−3^h, and Equation (16) is integrated with *Δx* = *Δy* = 4 *μ*m, and *Δt* = 1 *×* 10*^−3^* h. The remaining parameters are *N* = 15, *a* = 0.05/*μ*m, *f*_0_ = 1 *μ*m/h.

**Fig. 5.**
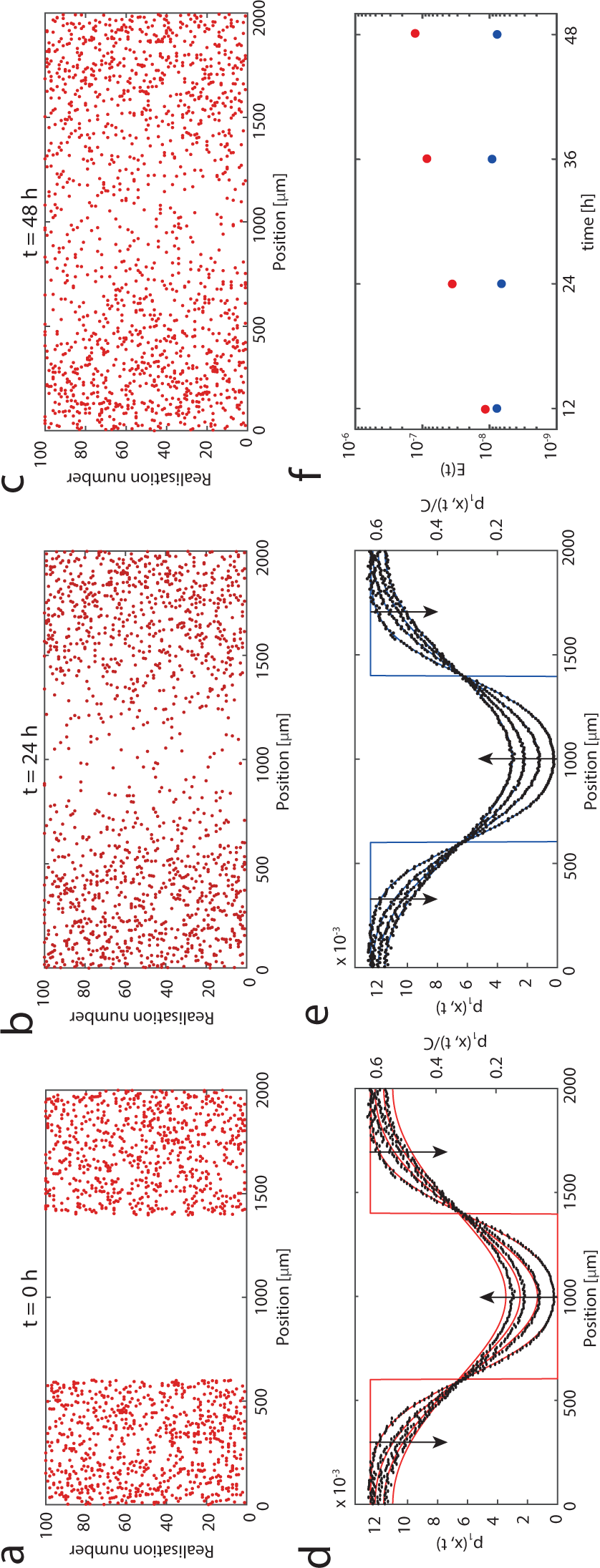
Comparison of the averaged agent density data from stochastic simulations and the solutions of the MFA- and KSA-based continuum models for a scratch assay with agents where the diameter grows relatively slowly, with *k* = 0.2 /h. Snapshots in (a)-(c) show 100 identically prepared realisations of the stochastic model, given by Equation (3), at *t* = 0,24 and 48 h, respectively. The initial density is given by Equation (17). Results in (d) and (e) show the averaged agent density profiles using an ensemble of 10^5^ identically prepared simulations (black dots) with binsize 10 *μ*m. The averaged results are compared with solutions of the MFA-based continuum model, Equation (13) (red lines), and the KSA-based continuum model, Equation (16) (blue lines). Profiles are given at *t* = 0, 12, 24, 36 and 48 h, with the arrows indicating the direction of increasing *t*. Results in (f) show the evolution of the mean squared errors (Equations (18) and (19)) for the MFA- and KSA-based continuum models. Equation (3) is integrated with *Δt* = 2 *×* 10*^−2^ h*, Equation (13) is integrated with *Δx* = 4 *μ*m and *Δt* = 1 *×* 10*^−3^* h, and Equation (16) is integrated with *Δx* = *Δy* = 4 *μ*m, and *Δt* = 1 *×* 10*^−3^* h. The remaining parameters are *N* = 15, *a* = 0.05/*μ*m, *f*_0_ = 0.1 *μ*m/h.

**Fig. 6.**
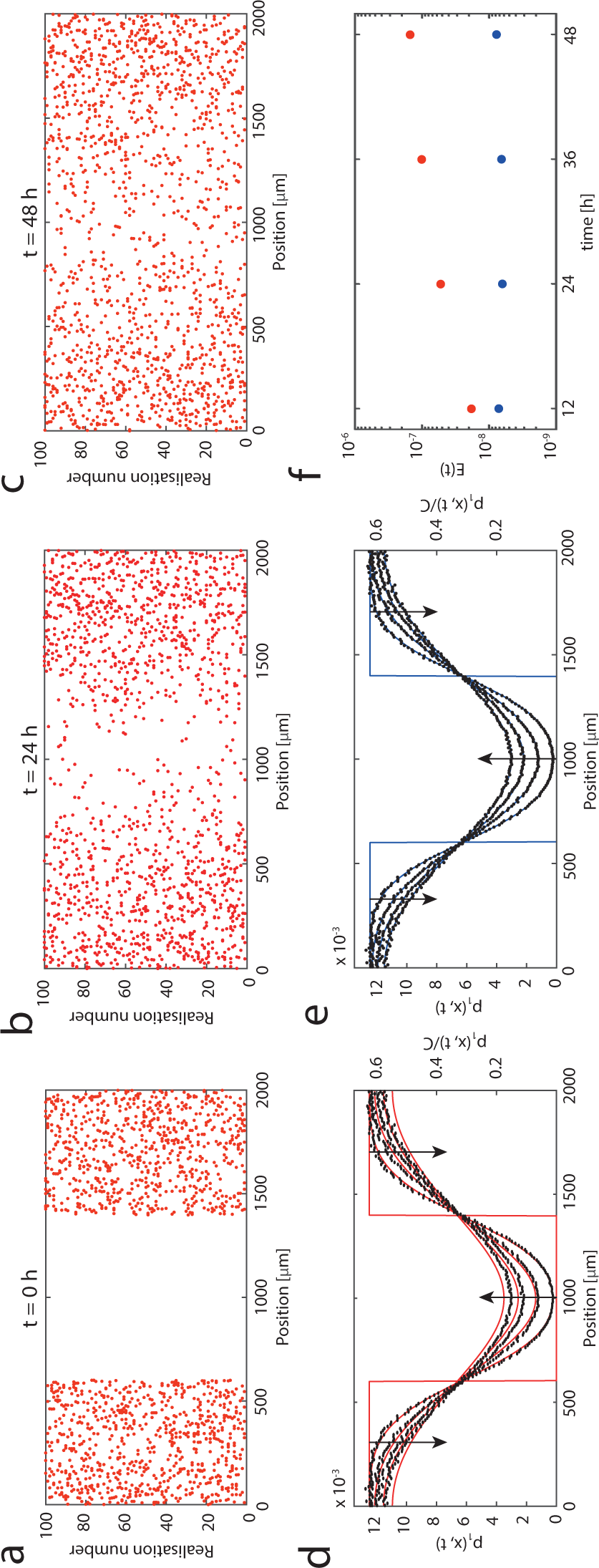
Comparison of the averaged agent density data from stochastic simulations and the solutions of the MFA- and KSA-based continuum models for a scratch assay with agents where the diameter grows faster, with *k* = 0.3 /h. Snapshots in (a)-(c) show 100 identically prepared realisations of the stochastic model, given by Equation (3), at *t* = 0, 24 and 48 h, respectively. The initial density is given by Equation (17). Results in (d) and (e) show the averaged agent density profiles using an ensemble of 10^5^ identically prepared simulations (black dots) with binsize 10 *μ*m. The averaged results are compared with solutions of the MFA-based continuum model, Equation (13) (red lines), and the KSA-based continuum model, Equation(16) (blue lines). Profiles are given at *t* = 0, 12, 24, 24, 36 and 48 h, with the arrows indicating the direction of increasing *t*. Results in (f) show the evolution of the mean squared errors (Equations (18) and (19)) for the MFA- and KSA-based continuum models. Equation (3) is integrated with *Δt* = 2 *×* 10*^−2^* h, Equation (13) is integrated with *Δx* = 4 *μ*m and *Δt* = 1 *×* 10*^−3^* h, and Equation (16) is integrated with *Δx = Δy* = 4 *μ*m, and *Δt* = 1 *×* 10*^−3^*h. The remaining parameters are *N* = 15, *a* = 0.05/*μ* m, *f*_0_ = 0.1 *μ*m/h.

## 4 Conclusions

In this work we present stochastic and continuum models of collective cell migration that can be applied to mimic scratch assays. In particular, we pay careful attention to allow for the case where the agents in the stochastic simulation change size dynamically. This feature can be important when we consider scratch assays with cells that are treated with Mitomycin-C to prevent proliferation. The stochastic model we present takes the form of a system of Langevin equations, and this framework can be used to describe the collective behaviour of a population of cells with constant size, or the collective behaviour of a population of cells with variable size. In addition to considering variable cell size, the stochastic framework describes random cell motility, crowding effects via a short range repulsive force, and cell-to-cell adhesive effects via longer range attraction. There is a crucial difference between two key parameter regimes that we consider: constant agent size, and dynamically increasing agent size. In the case when agent size remains constant, the average force acting on each individual agent remains approximately constant in regions of spatially uniform density. In contrast, increasing the size of individual agents leads to increased interactions between agents. Consequently, when we consider cases where the agent size increases dynamically, the MFA continuum description provides poorer match to the averaged discrete results at later times (Figure 5d, Figure 6d).

In addition to relying on repeated stochastic simulations, we also wish to develop continuum approximations of the stochastic model so that we can predict population-level and tissue-level data. To achieve this we first consider a continuum description based on the usual MFA that neglects correlations in the positions of agents. In this approach the position of any individual agent is treated as being independent of the positions of all other agents. The MFA-based model is relatively fast to simulate, as it takes only a few minutes to produce results depicted in Figure 4d, Figure 5d, and Figure 6d on a single desktop machine. While the MFA leads to a straightforward continuum model, the neglect of spatial correlations suggests that the approach might not always be valid, for example in situations where spatial structure and clustering develops. To overcome these potential limitations we also make use of a more advanced moment dynamics approach using the KSA. The KSA-based continuum description is more complicated to derive and much more numerically intense than the MFA-based approach, but it is attractive since it avoids inaccuracies of the MFA.

Generally, both the MFA- and KSA-based continuum models lead to reasonable predictions of the averaged stochastic results for the experimentally motivated problems that we describe here. In the case when the agent size remains constant, both the MFA- and KSA-based continuum models lead to an excellent match with the averaged data from the stochastic simulations. In this case, the simpler MFA-based continuum model is preferable to the computationally expensive KSA-based continuum model. However, in cases where agents increase in size, the KSA-based model outperforms the MFA-based model. This is due to the fact that agent growth increases agent-to-agent crowding effects, and these effects are incorporated in a relatively simplistic way in the MFA-based continuum model. Instead of simply concluding that the KSA-based model is always preferable to the MFA-based model, we acknowledge that the increased accuracy of the KSA-based approach comes at the cost of significantly increased computational expense. Specifically, it takes a couple of days to produce the results shown in Figure 4e, Figure 5e, and Figure 6e on High Performance Computing facilities without parallelizing techniques (QUT High Performance Computing, 2017). Therefore, we take a flexible view and present both continuum models. Furthermore, we acknowledge that the MFA-based approach will be preferable in some circumstances, whereas the KSA-based approach will be preferable in other circumstances.

There are many ways that the work presented here can be extended. For example, all cases presented here involve particular choices of functional forms for *δ*(*t*) and *Z*(*r, t*), yet many other choices are possible. Note that the stochastic algorithm described here, and the two continuum approximations are sufficiently flexible that other functional forms for *δ*(*t*) and *Z*(*r, t*) can be used directly in these frameworks, if required. We note that, here we consider the change in cell diameter since this is the simplest possible way that we can mimic an increase in cell size. However, alternative approaches are possible, such as modelling dynamic changes in cell volume. Our modelling approach can be used to mimic dynamic changes in volume by assuming that cells are spherical, and expressing the radius as a function of volume. Here we do not pursue this approach as the experimental images in Figure 1 provide little information about the three-dimensional shape of the cells, so we feel it is more natural to work with a simpler measure, namely the approximate diameter, *δ*(*t*). Another assumption that we make is that all agents in the population behave identically in that each agent has the same initial size and grows at the same rate. An interesting extension of this work would be to consider a heterogeneous population of cells that is made up of distinct subpopulations. Using the framework presented it would be possible to consider different subpopulations with different initial sizes, and to consider different subpopulations that grow at different rates. This kind of model could be described using a more complicated multi-species framework (Matsiaka et al. 2017). However, since this is the first time that a model of collective cell migration in a scratch assay that incorporates crowding effects and cell size dynamics has been explored, we leave this extension to the multi-species case for future consideration.

## Acknowledgements

We appreciate support from the Australian Research Council (FT130100148 and DP170100474). Ruth E Baker is a Royal Society Wolfson Research Merit Award holder and a Leverhulme Research Fellow. Computational resources are provided by the High Performance Computing and Research Support Group at QUT.

## References

1. Baker RE, Simpson MJ (2010). Correcting mean-field approximations for birth-death-movement processes. Physical Review E. 82: 041905.

2. Binder BJ, Simpson MJ (2016). Cell density and cell size dynamics during in vitro tissue growth experiments: Implications for mathematical models of collective cell behaviour. Applied Mathematical Modelling. 40: 3438-3446.

3. Binny RN, Plank MJ, James A (2015). Spatial moment dynamics for collective cell movement incorporating a neighbour-dependent directional bias. Journal of the Royal Society Interface 12: 20150228.

4. Binny RN, Haridas P, James A, Law R, Simpson MJ, Plank MJ (2016). Spatial structure arising from neighbour dependent bias in collective cell movement. PeerJ 4:e1689.

5. Binny RN, James A, Plank MJ (2016). Collective cell behaviour with neighbour-dependent proliferation, death and directional bias. Bulletin of Mathematical Biology. 78(11): 2277-2301.

6. Bolker B, Pacala SW (1997). Using moment equations to understand stochastically driven spatial pattern formation in ecological systems. Theoretical Population Biology. 52: 179-197.

7. Byrne H, Preziosi L (2003). Modelling solid tumour growth using the theory of mixtures. Mathematical Medicine and Biology. 20: 341-366.

8. Callaghan T, Khain E, Sander LM, Ziff RM (2006). A stochastic model for wound healing. Journal of Statistical Physics. 122(5): 909-924.

9. Codling EA, Plank MJ, Benhamou S (2008). Random walk models in biology. Journal of the Royal Society Interface. 5: 813-834.

10. Dyson L, Maini PK, Baker RE (2012). Macroscopic limits of individual-based models for motile cell populations with volume exclusion. Physical Review E. 86: 031903.

11. Edmondson R, Broglie JJ, Adcock AF, Yang L (2014). Three-dimensional cell culture systems and their applications in drug discovery and cell-based biosensors. ASSAY and Drug Development Technologies. 12(4): 207-218.

12. Galle J, Loeffler M, Drasdo D (2005). Modeling the effect of deregulated proliferation and apoptosis on the growth dynamics of epithelial cell populations in vitro. Biophysical Journal. 88: 62-75.

13. Glenn HL, Messner J, Meldrum DR (2016). A simple non-perturbing cell migration assay insensitive to proliferation effects. Scientific Reports. 6:31694.

14. Grada A, Otero-Vinas M, Prieto-Castrillo F, Obagi Z, Falanga V (2017). Research techniques made simple: analysis of collective cell migration using the wound healing assay. Journal of Investigative Dermatology. 137: e11-e16.

15. House T (2014). Algebraic moment closure for population dynamics on discrete structures. Bulletin of Mathematical Biology. 77: 646-659.

16. Jeon J, Quaranta V, Cummings PT (2010). An off-lattice hybrid discrete-continuum model of tumor growth and invasion. Biophysical Journal. 98: 37-47.

17. Jin W, Shah ET, Penington CJ, McCue SW, Chopin LK, Simpson MJ (2016). Reproducibility of scratch assays is affected by the initial degree of confluence: Experiments, modelling and model selection. Journal of Theoretical Biology. 390: 136-145.

18. Johnston ST, Shah ET, Chopin LK, McElwain DLS, Simpson MJ (2015). Estimating cell diffusivity and cell proliferation rate by interpreting IncuCyte ZOOMTM assay data using the Fisher-Kolmogorov model. BMC Systems Biology. 9:38.

19. Kaighn ME, Narayan KS, Ohnuki Y, Lechner JF, Jones LW (1979). Establishment and characterization of a human prostatic carcinoma cell line (PC-3). Investigative Urology. 17: 16-23.

20. Keeling MJ, Rand DA, Morris AJ (1997). Correlation models for childhood epidemics. Proceedings of the Royal Society of London B: Biological Sciences. 264: 1149-1156.

21. Khain E, Katakowski M, Hopkins S, Szalad A, Zheng X, Jiang F, Chopp M (2011). Collective behavior of brain tumor cells: The role of hypoxia. Physical Review E. 83: 031920

22. Kirkwood JG (1935). Statistical mechanics of fluid mixtures. Journal of Chemical Physics. 3(5): 300-313.

23. Levin SA, Whitfield M (1994). Patchiness in marine and terrestrial systems: from individuals to populations. Philosophical Transactions: Biological Sciences. 343: 99-103.

24. Liang CC, Park AY, Guan JL (2007). In vitro scratch assay: a convenient and inexpensive method for analysis of cell migration in vitro. Nature Protocols. 2: 329-333.

25. Maini PK, McElwain DL, Leavesley DI (2004). Traveling wave model to interpret a wound-healing cell migration assay for human peritoneal mesothelial cells. Tissue Engineering. 10 (3-4): 475-82.

26. Markham DC, Simpson MJ, Baker RE (2015). Choosing an appropriate modelling framework for analysing multispecies co-culture cell biology experiments. Bulletin of Mathematical Biology. 77: 713-734.

27. Matsiaka OM, Penington CJ, Baker RE, Simpson MJ (2017). Continuum approximations for lattice-free multi-species models of collective cell migration. Journal of Theoretical Biology. 422: 1-11.

28. Middleton AM, Fleck C, Grima R (2014). A continuum approximation to an off-lattice individual-cell based model of cell migration and adhesion. Journal of Theoretical Biology. 359: 220-232.

29. Murray PJ, Edwards CM, Tindall MJ, Maini PK (2012). Classifying general nonlinear force laws in cell-based models via the continuum limit. Physical Review E. 85: 021921.

30. Murrell DJ, Dieckmann U, Law R (2004). On moment closures for population dynamics in continuous space. Journal of Theoretical Biology. 229: 421-432.

31. Newman TJ, Grima R (2004). Many-body theory of chemotactic cell-cell interactions. Physical Review E. 70: 051916.

32. Nyegaard S, Christensen B, Rasmussen JT (2016). An optimized method for accurate quantification of cell migration using human small intestine cells. Metabolic Engineering Communications. 3: 76-83.

33. Penington CJ, Hughes BD, Landman KA (2011). Building macroscale models from microscale probabilistic models: A general probabilistic approach for nonlinear diffusion and multispecies phenomena. Physical Review E. 84: 041120.

34. Plank MJ, Law R (2015). Spatial point processes and moment dynamics in the life sciences: a parsimonious derivation and some extensions. Bulletin of Mathematical Biology. 77: 586-613.

35. Press WH, Flannery BP, Teukolsky SA, Vetterling WT (2007). Numerical recipes: The art of scientific computing (3rd ed.). Cambridge University Press. Cambridge. UK.

36. QUT High Performance Computing. https://www.student.qut.edu.au/technology/researchcomputing/high-performance-computing. (1 November 2017, date last accessed).

37. Riss T (2005). Selecting cell-based assays for drug discovery screening (2005). Cell Notes. 13: 16-21.

38. Sadeghi HM, Seitz B, Hayashi S, LaBree L, McDonnell PJ (1998). In vitro effects of Mitomycin-C on human keratocytes. Journal of Refractive Surgery. 14: 534-540.

39. Shah ET, Upadhyaya A, Philp LK, Tang T, Skalamera D, Gunter J, Nelson CC, Williams ED, Hollier BG (2016). Repositioning “old” drugs for new causes: identifying new inhibitors of prostate cancer cell migration and invasion. Clinical & Experimental Metastasis. 33: 385-399.

40. Sharkey KJ, Fernandez C, Morgan KL, Peeler E, Thrush M, Turnbull JF, Bowers RG (2006). Pair-level approximations to the spatio-temporal dynamics of epidemics on asymmetric contact networks. Journal of Mathematical Biology. 53: 61-85.

41. Sharkey KJ, Kiss IZ, Wilkinson RR, Simon PL (2015). Exact equations for SIR epidemics on tree graphs. Bulletin of Mathemtical Biology. 77: 614-645.

42. Sherratt JA, Murray JD (1990). Models of epidermal wound healing. Proceedings of the Royal Society of London B. 241: 29-36.

43. Simpson MJ (2009). Depth-averaging errors in reactive transport modelling. Water Resources Research. 45: W02505.

44. Simpson MJ, Treloar KK, Binder BJ, Haridas P, Manton KJ, Leavesley DI, McElwain DLS, Baker RE (2013). Quantifying the roles of cell motility and cell proliferation in a circular barrier assay. Journal of the Royal Society Interface. 10: 20130007.

45. Singer A (2004). Maximum entropy formulation of the Kirkwood superposition approximation. The Journal of Chemical Physics. 121: 3657-3666.

46. Smith S, Cianci C, Grima R (2017). Macromolecular crowding directs the motion of small molecules inside cells. Journal of the Royal Society Interface. 14: 20170047.

47. Steinberg MS (1996). Adhesion in development: An historical review. Developmental Biology. 180: 377-388.

48. Tersoff J (1987). New empirical approach for the structure and energy of covalent systems. Physical Review B. 37(12): 6991-7000.

49. Treloar KK, Simpson MJ, Haridas P, Manton KJ, Leavesley DI, McElwain DLS, Baker RE (2013). Multiple types of data are required to identify the mechanisms influencing the spatial expansion of melanoma cell colonies. BMC Systems Biology. 7: 137.

